# LutABC of *Veillonella parvula* Deacidifies *Streptococcus mutans* Biofilms and Improves Biofilm Health

**DOI:** 10.64898/2026.08.17.745241

**Authors:** Joseph M. Ferracciolo, Haya B. Eldana, Carla Sena, Lea Chami, Sara A. Abdulelah, Neil A. Patel, Eric S. Krukonis

**Author notes:** Correspondence: Dr. Eric S. Krukonis, Division of Integrated Biomedical Sciences, University of Detroit Mercy School of Dentistry, 2700 Martin Luther King Jr. Blvd., Room 444, Detroit, MI 48208.

## Abstract

*S. mutans* and *V. parvula* cooperate in dental plaque to assemble a healthy biofilm and are associated with increased caries risk. *S. mutans* produces lactic acid from carbohydrates resulting in a final biofilm pH∼4, while *V. parvula* metabolizes lactate to acetic and propionic acids resulting in pH∼5. This process results in healthier biofilms that still generate a pH capable of demineralizing tooth surfaces (pH<5.5). The purpose of this study was to identify *V. parvula* genes required for deacidification of *S. mutans* biofilms and determine whether the ability of *V. parvula* to deacidify *S. mutans* biofilms correlates with enhanced biofilm health. Using transposon mutagenesis in *V. parvula* we identified several genes required for deacidification of *S. mutans* biofilms. These included numerous *V. parvula* transposon mutations in the previously unstudied *lutABC* lactate utilization operon. To assess biofilm health, *S. mutans* in the presence of various *V. parvula* mutants were stained with a LIVE/DEAD stain and imaged by fluorescence microscopy. An intact *lutABC* operon was required to enhance biofilm health, as demonstrated by plasmid-based complementation of a *lutB* transposon mutant. Transposon insertions in other loci unrelated to deacidification had no impact on biofilm health. Addition of HEPES buffer at the time of *S. mutans* biofilm assembly prevented full acidification of the biofilm and resulted in improved biofilm health, even without the addition of *V. parvula*. Finally, we found *V. parvula* can use either nitrate or fumarate as a final ETC electron acceptor during lactate utilization. In all, we found the *lutABC* lactate utilization operon of *V. parvula* is critical for the ability of *V. parvula* to deacidify *S. mutans* biofilms and promote biofilm health. Interfering with this pathway would interrupt the mutually beneficial relationship between *S. mutans* and *V. parvula* that leads to their co-association in caries, root caries, and early childhood caries.

## Introduction

While *S. mutans* is often considered the primary etiological agent of caries in humans (Fitzgerald and Keyes, 1960; Loesche et al., 1975), several members of the oral microbiota can also contribute to caries development (Chen et al., 2015; Fure et al., 1987; Gross et al., 2012; Preza et al., 2008). *V. parvula* is associated with caries, root caries and early childhood caries and its prevalence is correlated with the levels of *S. mutans* present in dental plaque (Abram et al., 2022; Chen et al., 2015; Gross et al., 2012). Since *V. parvula* utilizes lactate as a preferred carbon source it was hypothesized that it might reduce caries risk by reducing the levels of lactic acid in plaque (Mikx and Van der Hoeven, 1975). However, the by-products of lactate metabolism, acetic acid and propionic acid result in a biofilm pH=5, still below the critical pH=5.5 for enamel demineralization (Fig. 1, (Abram et al., 2022; Dawes, 2003)).

**Figure 1:**
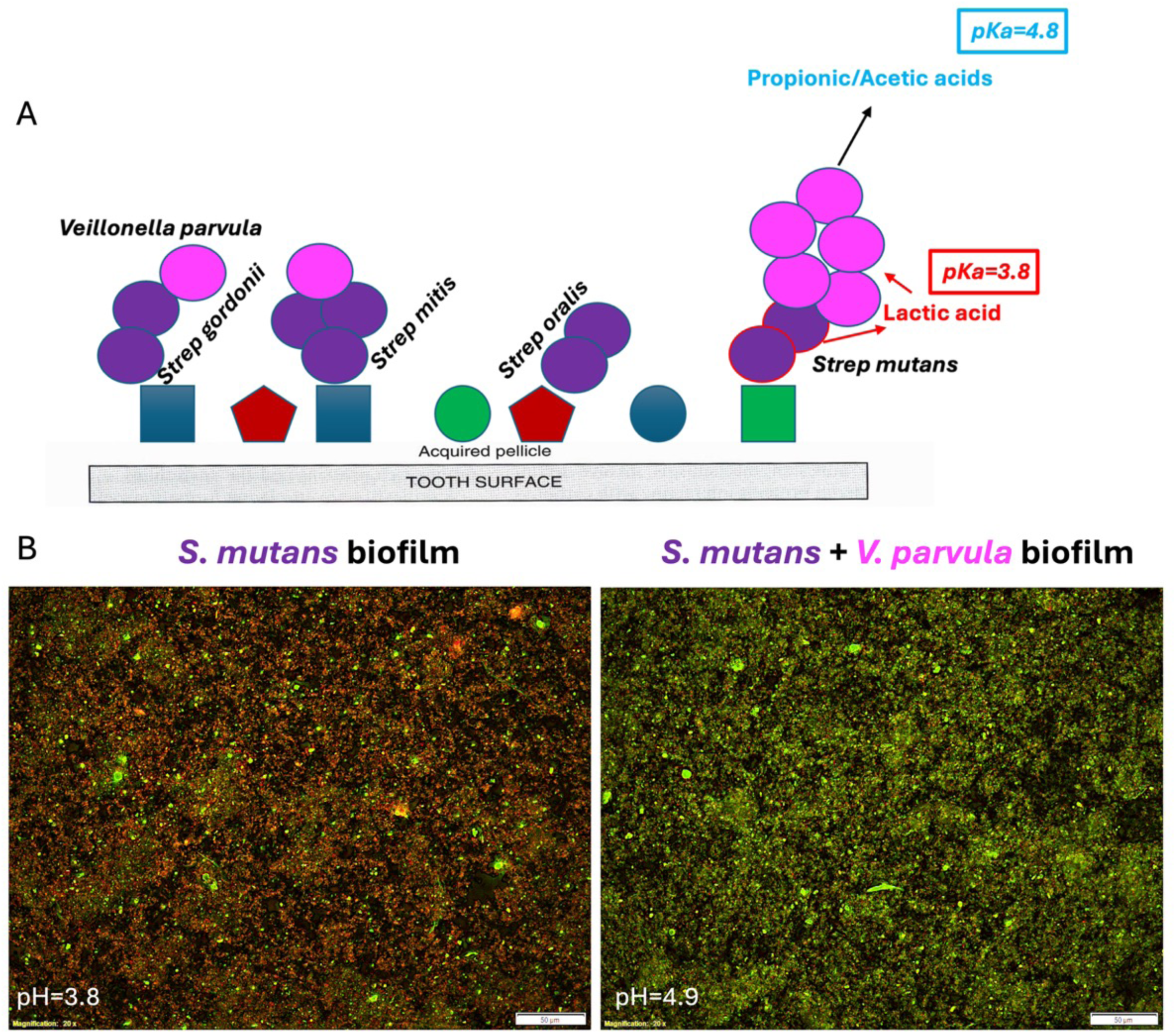
Model illustrating how *V. parvula* leads to deacidification of *S. mutans* biofilms (A). 24-hour biofilms of *S. mutans* alone and *S. mutans* (Sm) + *V. parvula* (Vp) were assessed for final pH of the biofilm supernatant and biofilm health via LIVE/DEAD staining (B).

In addition to deacidification of biofilms by *V. parvula*, *V. parvula* also enhances the health of 24 hr *S. mutans* biofilms in artificial saliva with sucrose, a medium reflective of the composition of human saliva (Abram et al., 2022; Roger et al., 2011). How *V. parvu*la contributes to the health of mixed biofilms is not fully understood, but we hypothesize reduced acidity plays a role. Alternatively, reduction in lactate may lead to enhanced biofilm health as has been described with the relationship between *Corynebacterium matruchotii* and *Streptococcus mitis* (Almeida et al., 2023).

In this study we employed a transposon mutagenesis strategy recently developed for *V. parvula* (Bechon et al., 2020) and sought to identify *V. parvula* genes required for deacidification of *S. mutans* biofilms. We performed 96-well biofilm assays in the presence of the pH-sensitive dye, bromocresol green, that is green at pH=5 and yellow at pH≤4.0 (Fig. 2A). Using this approach, we identified numerous transposon insertions in the previously unstudied lactate utilization operon of *V. parvula*: *lutABC,* as well as a locus encoding an ABC transporter substrate-binding protein, Vp_0785. Mutants from this screen were then assessed for their ability to improve the health of mixed biofilms.

**Figure 2:**
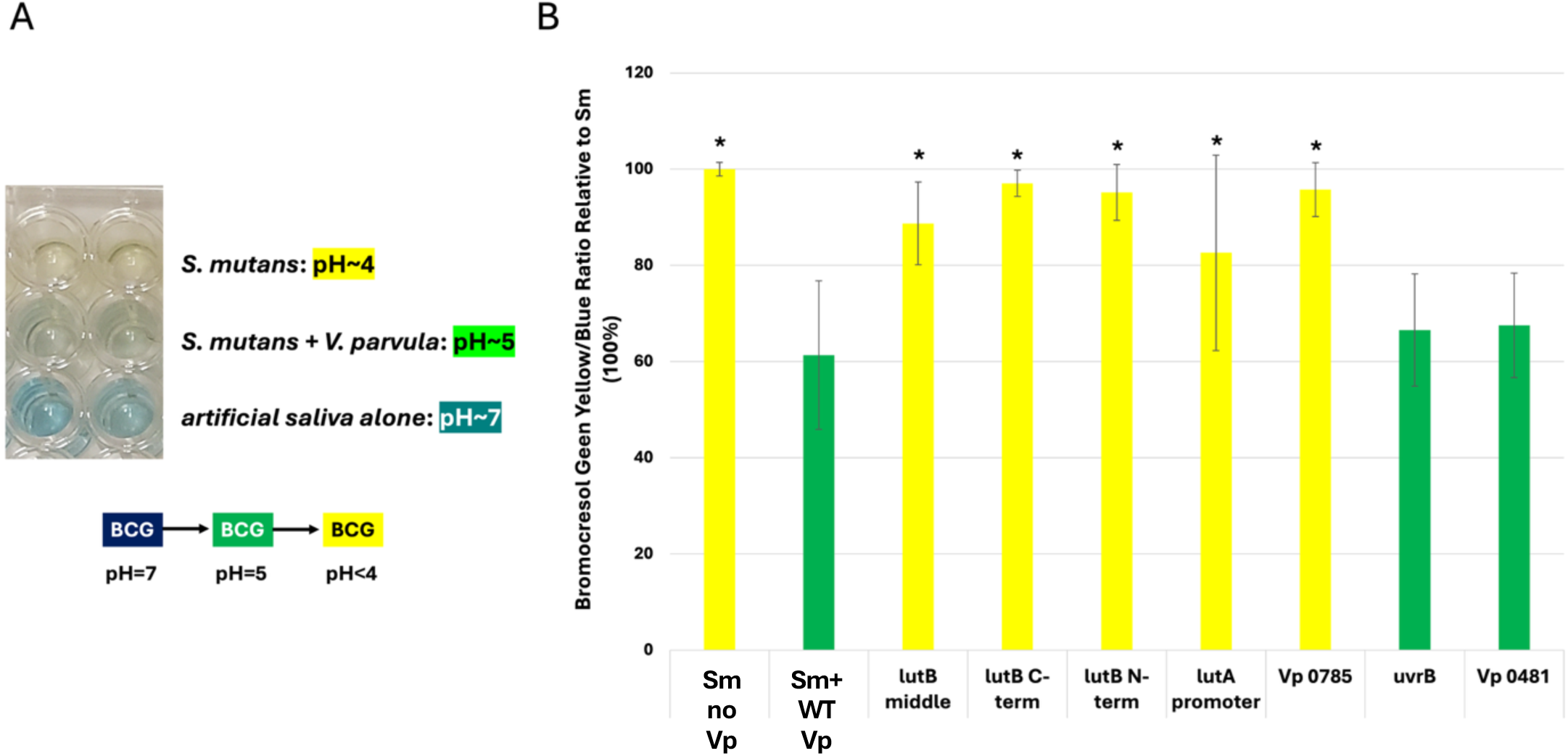
Screening for deacidification mutants of *V. parvula* using the pH-sensitive dye, bromocresol green (A). 24-hour biofilms of *S. mutans* alone and *S. mutans* + *V. parvula* or various *V. parvula* transposon mutants co-cultured with *S. mutans* were assessed for final pH of the biofilm supernatant using the colorimetric dye bromocresol green (B). The ratio of yellow:blue is shown n=2-4 (triplicate measurements each experiment). * p<0.05 by Students’ T-test.

## Methods

### Bacterial strains and growth conditions

*S. mutans* (ATCC 25175) and *V. parvula* (ATCC 10790) were grown overnight in BHI or Brucella broth + 1μg/ml vitamin K and 5μg/ml hemin, respectively. *S. mutans* was grown at 37°C 5% CO_2_ without shaking. *V. parvula* was grown at 37°C anaerobically without shaking. Alternatively, *V. parvula* was grown SK medium with 0.6% sodium lactate (Sigma cat#L4263; (Knapp et al., 2017)).

### Transposon mutant library assembly

A Mariner-based transposon mutant library was assembled in *V. parvula* according to the methods of Bechon *et al*, (Bechon et al., 2020). Briefly a chloramphenicol-resistant plasmid pRPF215 carrying a transposon encoding erythromycin resistance was introduced into *V. parvula* via electroporation (Dembek et al., 2015). The transposase was then induced with 0.1μg/ml anhydrous tetracycline (aTc) to induce transposon insertion onto the chromosome as well as prevent plasmid replication. *V. parvula* was grown for 4 hours or overnight in BHI medium in the presence of 0.1% L-cysteine, 0.6% lactate and 0.1μg/ml aTc. Cells were then plated on BHI agar with 200 μg/ml erythromycin, 0.1% L-cysteine, 0.6% lactate and 0.1μg/ml aTc and colonies were screened for loss of the plasmid based on loss of chloramphenicol resistance. Colonies were then screened directly for the ability to deacidify *S. mutans* biofilms and were stored as glycerol stocks.

### Biofilm assembly and pH measurements

Overnight cultures of *S. mutans* (Sm) and *V. parvula* (Vp) were pelleted and washed once with 10ml PBS, then repelleted and resuspended at an OD_600_=1.0 in artificial saliva (Roger et al., 2011) + 2mg/ml porcine mucin (Sigma cat#M1778) + 0.5% sucrose. 24-well plates or 4-well chamber slides were inoculated with 250μl Sm +/- 250μl Vp and the total well volume was brought to 1ml with artificial saliva with sucrose. Plates and slides were grown overnight at 37°C anaerobically and then processed for pH measurements and LIVE/DEAD staining.

### Use of bromocresol green dye

To screen for *V. parvula* transposon mutants unable to deacidify *S. mutans* biofilms, 96-well biofilms were assembled as described above in artificial saliva with 0.5% sucrose except the final volume was 100μl rather than 1ml and 0.3% bromocresol green (Sigma cat#114359) was included in the biofilm mixtures. Levels of acidification were visualized (Fig. 2A) and quantified by plotting the yellow absorbance (ABS=445nm)/blue absorbance (ABS=620nm), after subtracting the yellow/blue ratio of artificial saliva alone.

### Biofilm health assessment via ffuorescence microscopy using LIVE/DEAD staining

Following overnight growth, biofilms containing Sm or Sm + Vp were washed once with 1ml PBS and then a mixture of 2μg/ml SYTO9 and 20μg/ml propidium iodide in 250μl PBS were added to assess cell permeability. Cells with leaky/damaged membranes stain RED, while those with intact membranes stain GREEN. Biofilm were incubated for 10 minutes with the dyes at room temperature in the dark and then imaged on an Olympus BX63 fluorescence microscope and subjected to deconvolution imaging. At least two fields/well were imaged to ensure images were representative of the field. In some cases, parallel wells were processed for Gram straining to visualize *S. mutans* (Gram+ cocci) and *V. parvula* (Gram-cocci). In some cases, various concentrations of HEPES buffer pH=7.0, sodium lactate (Sigma cat#L4263), or lactic acid (Sigma cat#W261106) were added to the biofilm wells. When DNAse was used to determine if the source of propidium iodide straining (RED) was extracellular DNA, 5 units (2.5μl) of DNAse (New England Biolabs cat#M0303S) in 250μl 1X DNAse buffer were added to wells for 30 min at 37°C, after rinsing the wells once with 1ml PBS. Wells were then washed again twice with 1ml PBS, prior to imaging.

### lutABC mutant complementation

To ensure the phenotypes associated with *lutABC* transposon mutants were due to alterations in the *lutABC* locus, we assembled a complementing plasmid based on the *E.coli*/*V. parvula* shuttle vector pCF1135 (Goetting-Minesky et al., 2024). *lutABC* was amplified from the *V. parvula* 10790 genome using the NotI containing primers, (forward primer, 5’ cgcgGCGGCCGCgaaaagggctcctttcaaactatac 3’; reverse primer 5’ cgcgGCGGCCGCaatttttgataacatttaatgtattc 3’). The NotI-digested PCR product was ligated into NotI-digested pSK-Bluescript. Next the BamHI site in *lutC* was mutated while retaining the amino acid sequence with primers (forward primer, 5’ cttccaatgggaCccagctaaag 3’; reverse primer 5’ ctttagctggGtcccattggaag 3’). Finally, *lutABC* was reamplified from pSK-*lutABC* BamHI^-^ with BamHI containing primers (forward primer, 5’ cgcgGGATCCgaaaagggctcctttcaaactatac 3’; reverse primer 5’ cgcgGGATCCaatttttgataacatttaatgtattc 3’), digested with BamHI and ligated into BamHI cut pCF1135. Candidate colonies were sequenced and wild-type *lutABC* clones were introduced into a *V. parvula* transposon mutant in *lutB* for complementation via electroporation (Goetting-Minesky et al., 2024).

## Results

### V. parvula reduces the acidity of S. mutans biofilms and improves biofilm health

To demonstrate the effect of *V. parvula* on *S. mutans* biofilms, chamber slide biofilms were assembled, and pH and biofilm health were measured after 24 hrs. As previously reported (Abram et al., 2022), in artificial saliva +0.5% sucrose, dense *S. mutans* biofilms reached a pH of ∼4 and become unhealthy over 24 hours and stain with propidium iodide, indicating membrane damage (Fig. 1B). Inclusion of *V. parvula,* at an equivalent concentration, results in a much healthier biofilm at 24 hours. Notably, the pH of the resulting biofilm is ∼1 pH unit higher (pH∼5, 10-fold reduction in H^+^ ions)) when *V. parvula* is present (Fig. 1B). This pH shift is due to the ability of *V. parvula* to metabolize lactic acid (lactate, pKa=3.8) into the two weaker organic acids, propionic acid and acetic acid (pKa’s 4.9 and 4.8, respectively; (Ng and Hamilton, 1971)).

### Numerous V. parvula lutABC transposon mutants fail to prevent acidification of S. mutans biofilms

To identify *V. parvula* genes required for deacidification of *S. mutans* biofilms, we assembled a transposon library in *V. parvula* and assessed individual mutants for the ability to deacidify *S. mutans* biofilms using the pH-sensitive dye bromocresol green (BCG, Fig. 2A).

Three pools of erythromycin-resistant *V. parvula* mutants (due to expression of the transposon-encoded *ermB* gene, (Dembek et al., 2015)) were assembled and individual Erm-resistant mutants were inoculated into 96-well plates in triplicate with *S. mutans* in the presence of artificial saliva with 0.5% sucrose and 0.3% BCG. Wells containing *S. mutans* alone became yellow overnight as the pH approached pH=4, while wells with wild-type *V. parvula* and *S. mutans* were green, indicating a pH∼5 (Fig. 2A). Several *V. parvula* transposon mutants lost the ability to deacidify *S. mutans* biofilms, these included several mutants within the previously uncharacterized *lutABC* locus of *V. parvula* (Fig. 2B). This locus in other organisms is required for lactate utilization (Almeida et al., 2023; Chai et al., 2009; Sinha et al., 2024; Thomas et al., 2011), and plays a similar function in *V. parvula*. Transposon insertion sites were determined via PCR and sequencing of the transposon junction. Three unique insertion-site mutants were found in *lutB* and one was found in the *lutA* promoter (Figure 2B). One additional Tn mutant was found in locus Vp_0785, an ABC-transporter substrate-binding protein homolog. As controls, we show two *V. parvula* Tn insertion mutants that had no effect on the ability of *V. parvula* to deacidify *S. mutans* biofilms; *uvrB* and Vp_0481.

### lutABC mutants of V. parvula no longer improve S. mutans biofilm health

Previous studies suggested a link between acidification of dense *S. mutans* biofilms and membrane permeability (Fig. 1B, (Abram et al., 2022)). Given the role of the *lutABC* operon of *V. parvula* in *S. mutans* biofilm deacidification, we determined whether *lutABC* mutants were also unable to improve *S. mutans* biofilm health. Biofilms containing *S. mutans* alone, *S. mutans* with wild-type *V. parvula*, or *S. mutans* with a transposon mutant were grown for 24 hours. Addition of wild-type *V. parvula* raised the biofilm pH from pH=4 to pH=5 and the biofilm was healthier (green). However, when a *V. parvula lutB* or *lutA* promoter transposon mutant was co-cultured with *S. mutans*, there was no increase in pH and no health benefit (Fig. 3A). This was not due to the inability of *V. parvula* to become established in the biofilm as Gram staining showed equivalent amounts of *S. mutans* (Gram+) and *V. parvula* (Gram-) in the final biofilm (Fig. 3B). Transposon insertion in a *V. parvula* gene unrelated to deacidification, *uvrB*, had no effect on the health benefits of *V. parvula*, ruling out an effect of the transposon itself. Reintroduction of *lutABC* on a complementing plasmid (pCF1135-*lutABC*) resulted in restoration of both deacidification and biofilm health improvements to a *lutB* transposon mutant, confirming the defects were due to the transposon insertion in *lutB* (Fig. 3A).

**Figure 3:**
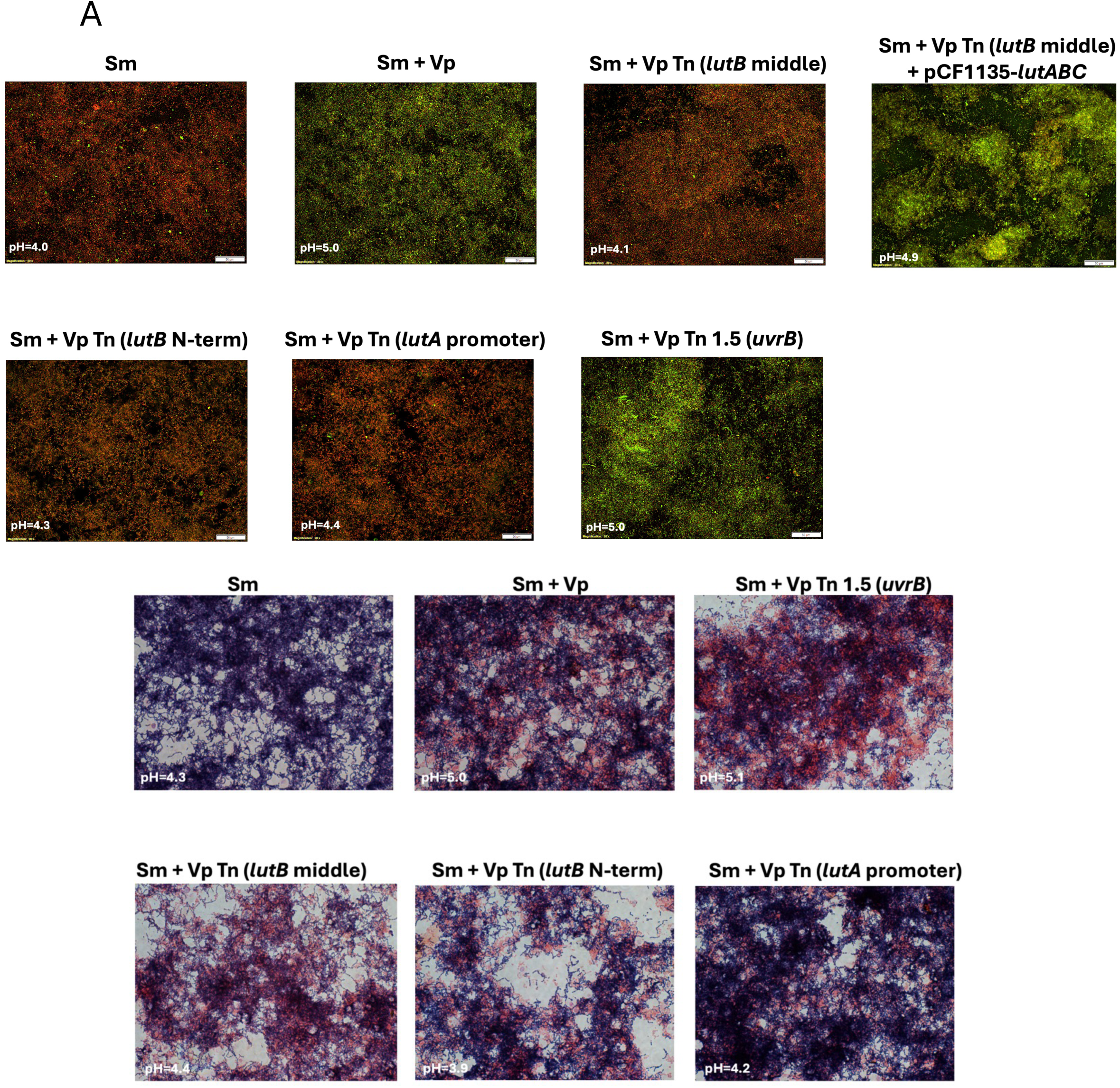
24-hour biofilms of *S. mutans* alone and *S. mutans* + *V. parvula* or various *V. parvula* mutants were assessed for final pH of the biofilm supernatant and biofilm health via LIVE/DEAD staining (A). pCF1135-lutABC is a complementing plasmid carrying the entire *lutABC* locus. Gram staining demonstrated both *S. mutans* and *V. parvula* are present in the final biofilm (B). Shown are representative images and typical pH results from at least 2 (Gram Stain)-3 (LIVE DEAD imaging) independent experiments.

We also investigated the possibility that the propidium iodine staining we see with *S. mutans* biofilms in the absence of *V. parvula* was due to extracellular DNA (eDNA, (Serrage et al., 2021)). We found treatment of the biofilms with DNAse prior to LIVE/DEAD staining resulted in no change in propidium iodine staining (Fig. 4).

**Figure 4:**
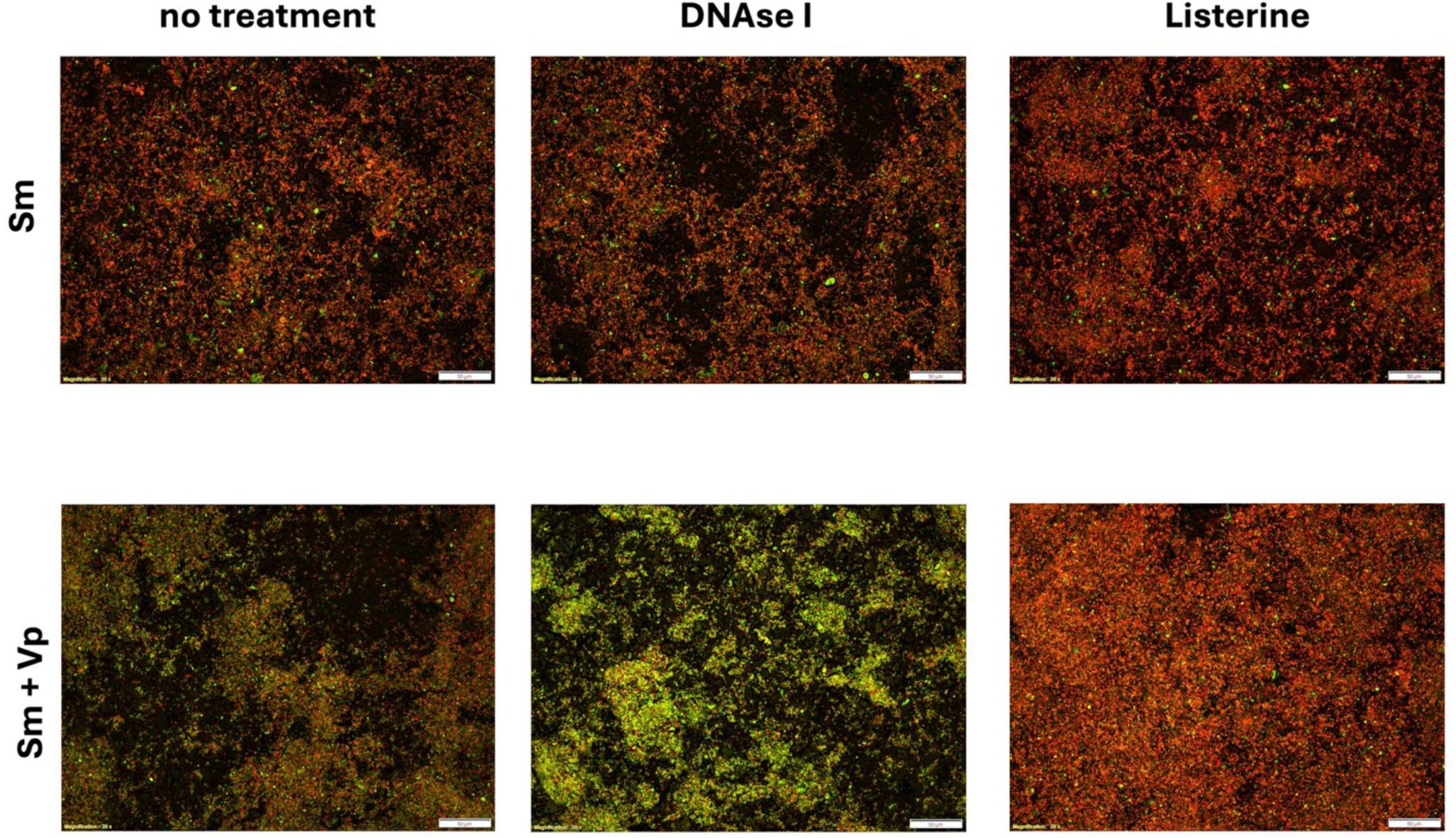
Propidium iodide staining of unhealthy *S. mutans* biofilms is not due to accessible extracellular DNA (eDNA). 24-hour biofilms of *S. mutans* or *S. mutans* + *V. parvula* were treated prior to LIVE/DEAD staining with 5 units of DNAse for 30 minutes. DNAse treatment had no impact on propidium iodide staining.

### Addition of buffering capacity to artificial saliva improves S. mutans biofilm health

One possible explanation for the membrane damaged observed with dense *S. mutans* biofilms is that prolonged exposure to a low pH environment (∼pH=4) eventually results in cell damage. To test this hypothesis, *S. mutans* biofilms were assembled as described above, but with increasing concentrations of HEPES buffer at pH=7. Addition of 100mM or 200mM HEPES resulted in healthier biofilms (green) with a final pH of ∼pH=5 and pH=6, respectively (Fig. 5A).

**Figure 5:**
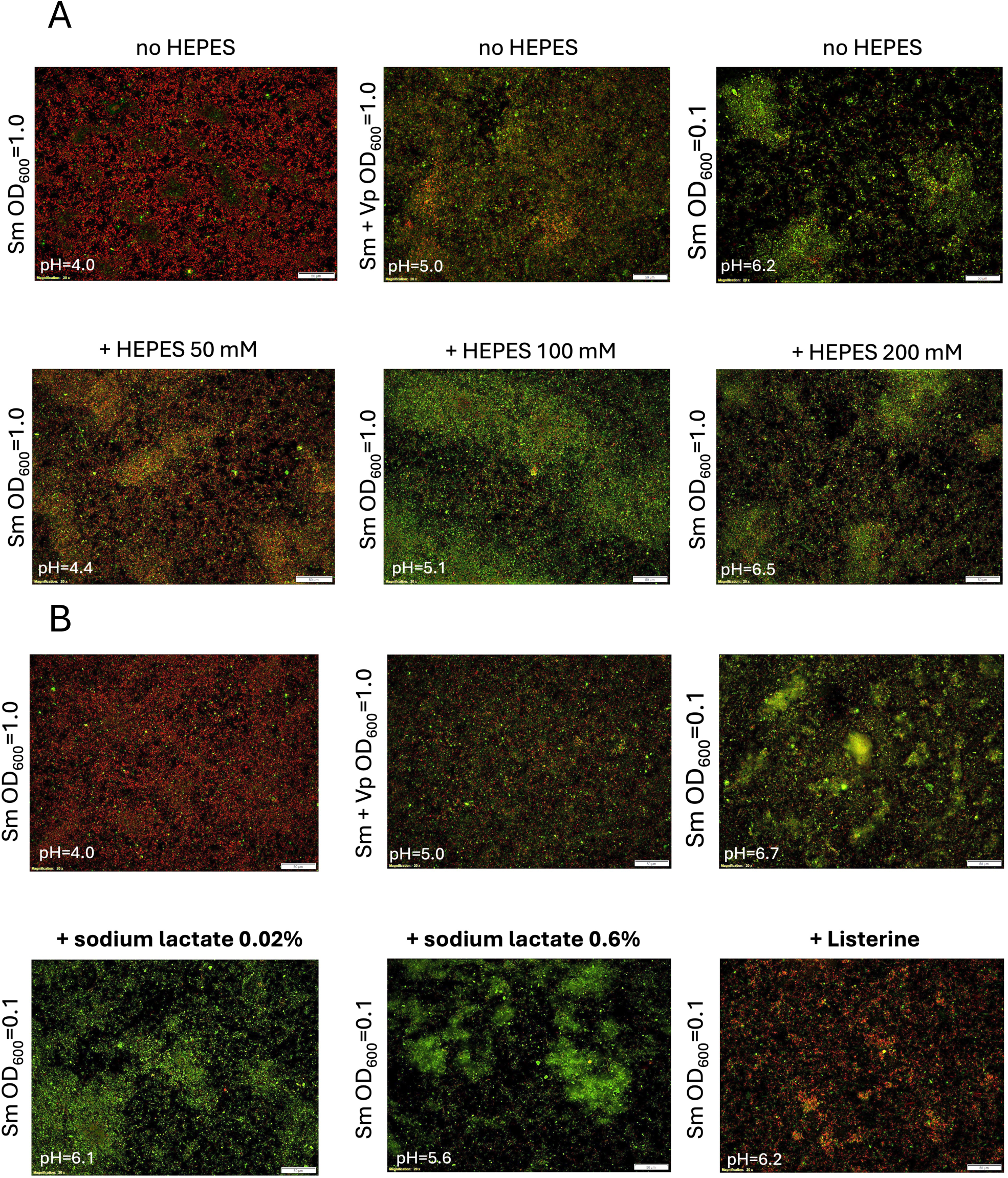
Buffering *S. mutans* biofilms with HEPES pH=7 mimics the deacidification and biofilm health benefits of *V. parvula*. Dense biofilms (OD_600_=1.0) were grown overnight in the presence or absence of increasing concentration of HEPES buffer (A). During biofilm assembly, 0.25ml of OD_600_=1.0 Sm was used in a total of 1ml. Thus, the starting OD_600_ was 0.25. The addition of sodium lactate did not impact biofilm health. Normally healthy moderately dense biofilms (OD_600_=0.1) were grown overnight in the presence of 0%, 0.02% or 0.6% sodium lactate (B).

An alternative potential source of toxicity to *S. mutans* in 24-hour biofilms is excess lactate (Almeida et al., 2023). Thus, we determined whether adding sodium lactate to low density *S. mutans* biofilms, prior to the typical time when lactic acid would build up to high levels in biofilms, could result in unhealthy *S. mutans* biofilms. Addition of either 0.02% lactate (the level in 24-hour *S. mutans* biofilms, ((Abram et al., 2022)) or 0.6% sodium lactate (the concentration of typically used for growth of *V. parvula* in culture) had no detrimental effect on 24-hour *S. mutans* biofilms initiated at a moderate density (0.25ml of OD_600_=0.1) as assessed by LIVE/DEAD staining (Fig. 5B). Our typical high-density *S. mutans* biofilms are established using 0.25ml of OD_600_=1.0. Treatment of biofilms with 100% Listerine Naturals prior to LIVE/DEAD imaging resulted in damaged cells, as indicated by red staining (Fig. 5B). The addition of lactic acid (rather than sodium lactate) was harder to assess for effects on biofilm health since at concentration as low as 0.08% lactic acid, no biofilms formed on the plate, and no LIVE/DEAD staining was possible (data not shown).

### Nitrate present in artificial saliva is required for lactate utilization by *V. parvula*

Under anaerobic growth conditions required for *V. parvula*, the final electron acceptor in the electron transport chain is often nitrate. To test whether *V. parvula* relies upon NaNO_3_ present in our artificial saliva formulation for *lutABC* effects, we made artificial saliva lacking NaNO_3_. Growth of *S. mutans* biofilms in artificial saliva-N0_3_ resulted in no deacidification by *V. parvula* (Fig. 6A) and no health benefits (Fig. 6B). Re-addition of 4.3mM NaNO_3_ restored deacidification and improved biofilm health (Fig. 6). Addition of 5mM fumarate as a substitute for NO_3_ gave partial deacidification and improved health of *S. mutans* biofilms in the presence of *V. parvula* (Fig. 6). Fumarate is an alternative electron acceptor under anaerobic conditions with less reduction potential than NO_3_ as an electron acceptor (Gennis, 1996).

**Figure 6:**
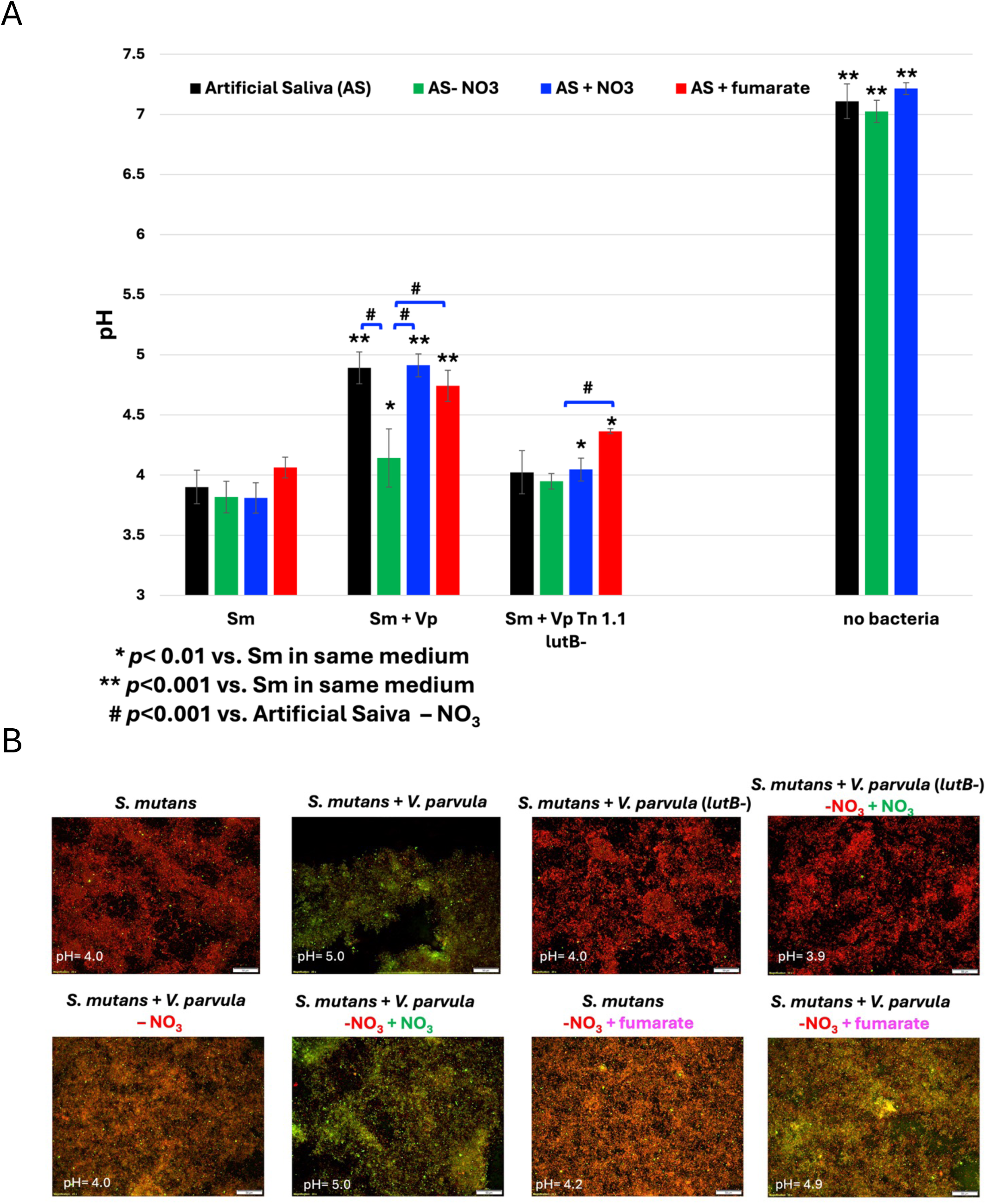
Removal of sodium nitrate from artificial saliva prevents *V. parvula* deacidification and improved health of *S. mutans* biofilms. pH measurements were performed on 24 hr biofilms in artificial saliva (AS), artificial saliva without sodium nitrate (AS-NO3), artificial saliva without sodium nitrate with 4.3mM sodium nitrate added back (AS+NO3), or artificial saliva without sodium nitrate with 5mM fumarate added back (AS + fumarate). A *lutB*-mutant was unable to respond to nitrate (A). 24 hr biofilms were grown in artificial saliva +/- 4.3mM sodium nitrate or 5mM fumarate and LIVE/DEAD imaging was performed. A *lutB*-mutant was included as a *V. parvula* mutant unable to respond to nitrate (B). Statistical analysis by the students’ T-test.

## Discussion

In this study we identified a pathway that enables *V. parvula* to deacidify *S. mutans* biofilms and improve biofilm health, the previously unstudied *V. parvula* lactate utilization system, LutABC.

We isolated five distinct transposon insertions (three in *lutB*, one in the *lutA* promoter, one in Vp_0785) that prevented *V. parvula* from deacidifying *S. mutans* biofilms by metabolizing lactate to propionate and acetate (Fig. 2B, (Ng and Hamilton, 1971)). Mutants affecting the *lutABC* operon also failed to improve the health of dense *S. mutans* biofilms (Fig. 3A). Supporting the hypothesis that an important role for *V. parvula* is deacidification of *S. mutans* biofilms, we found addition of HEPES buffer (pH=7) to *S. mutans* biofilms could also prevent acidification of the biofilm to pH=4, and this improved biofilm health (Fig. 5A). While one interpretation of these results is that *S. mutans* becomes sensitive to extended exposure to a low pH environment (∼pH=4), it is also possible that low pH environments trigger expression of genes in *S. mutans* that affect cell viability. While *S. mutans* biofilms stain RED in our LIVE/DEAD assays, they maintain 30-50% viability relative to *S. mutans* + *V. parvula* biofilms. Thus, while unhealthy, these *S. mutans* biofilms are still viable (Appendix Fig. 1). Alternatively, following a 10-minute treatment with Listerine Naturals no viable bacteria are recovered. While not the focus of this paper, future studies will explore the function of the protein encoded by Vp_0785. It encodes a putative ABC transporter substrate-binding protein, and its role may affect lactate import into *V. parvula* although it is not homologous to other lactate permeases associated with *lutABC* systems (*lctP*, (Thomas et al., 2011)).

The *lutABC* system was first described as playing a role in biofilm formation in *Bacillus subtilis* (Chai et al., 2009) and as a novel lactate dehydrogenase (LDH) in *Shewanella oneidensis* (Pinchuk et al., 2009). It has also been shown to be critical for *Campylobacter jejuni* to utilize host intestinal lactate and establish a productive infection in ferrets, a model mimicking human infection (Sinha et al., 2024; Thomas et al., 2011). In the oral cavity, the *lutABC* system of *Corynebacterium matruchotii* is important to support the health of *Streptococcus mitis* in mixed biofilms associated with oral health, but in this case, accumulating lactate appears to be the source of *S. mitis* toxicity (Almeida et al., 2023). Our results indicate the *V. parvula* LutABC system results in decreased acidity of biofilms accumulating in the oral cavity. Based on homology predictions LutA, LutB, and LutC are iron-sulfur cluster containing proteins that participate in oxidizing lactate to pyruvate in *V. parvula* and sequentially transfer elections via an electron transport chain (Hwang et al., 2013). Since *V. parvula* is an obligate anaerobe, the final electron acceptor for this process is likely nitrate, with nitrate reductase leading to the conversion of nitrate to nitrite and the generation of ATP for *V. parvula* (Fig. 7). We have shown nitrate or fumarate can serve as the final electron acceptor for the *V. parvula* LutABC system (Fig. 6).

**Figure 7:**
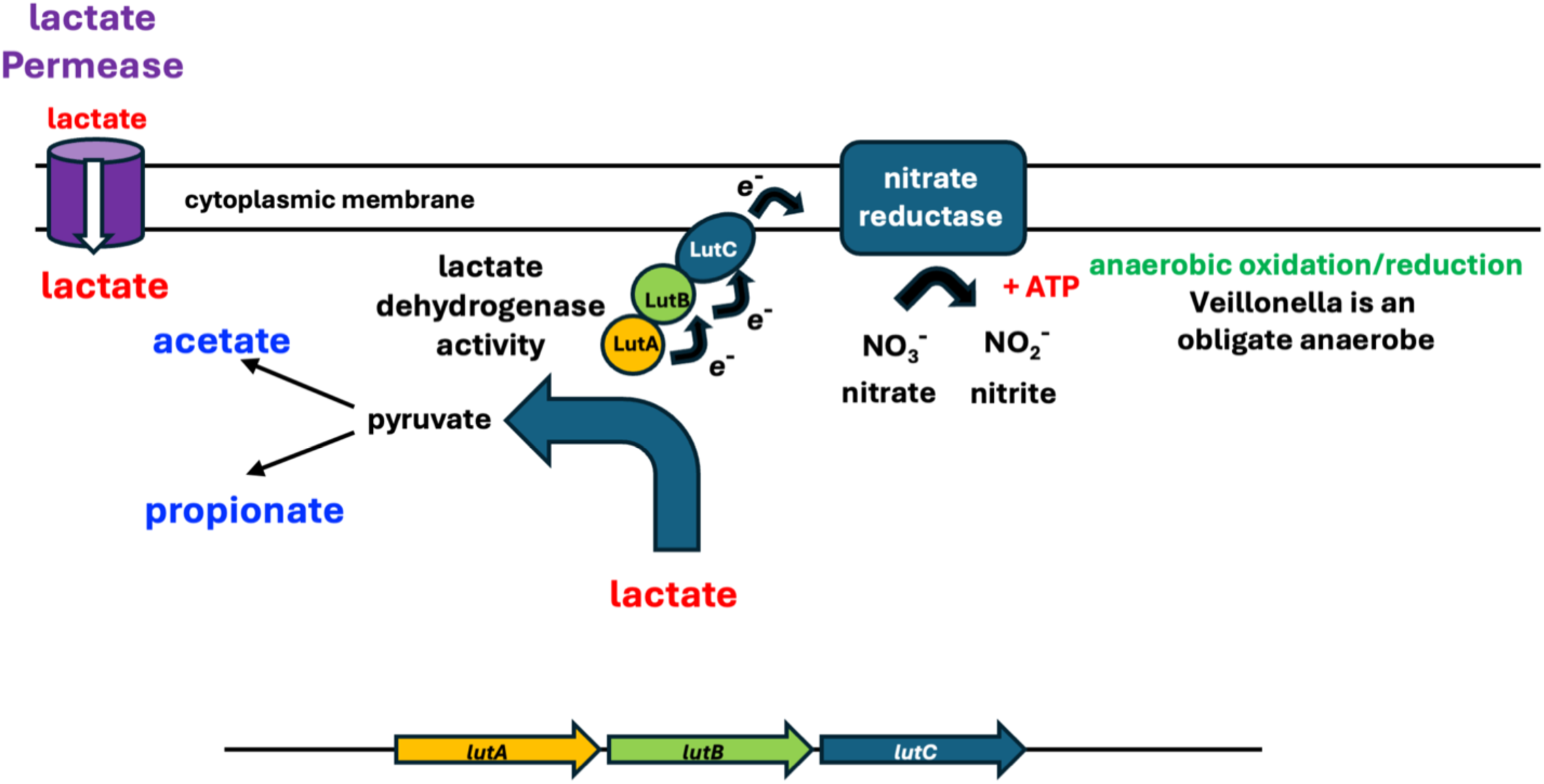
Model of LutABC Complex Function. The LutABC complex is proposed to constitute an electron transport chain using the conversion of nitrate to nitrite under anaerobic conditions to generate ATP upon lactate utilization. Figure modeled on Chai *et al* 2009.

It is important to note the final pH of dual species *S. mutans*/*V. parvula* biofilms in our artificial saliva model system is ∼pH=5, still below the critical pH for enamel demineralization (Dawes, 2003). This would explain why numerous studies have found an association of higher levels of V*eillonella species* on carious lesions in humans (Aas et al., 2008; Abram et al., 2022; Becker et al., 2002; Gross et al., 2012). Our findings also suggest these increased levels of *V. parvul*a are not just an indication of *V. parvula* benefitting from lactate-producing bacteria in dental plaque, but that the presence of *V. parvula* also provides some benefit to other members of plaque microbial community, like *S. mutans* and potentially other streptococci. Two recent studies showed combining *V. parvula* and *S. mutans* in a rat enamel or root caries models leads to more severe caries than *S. mutans* alone (Li et al., 2024; Wei et al., 2024). Future studies will determine the role of the *V. parvula* LutABC system in an animal model of caries.

## Acknowledgements

This work was funded by the University of Detroit Mercy School of Dentistry FRG #UDMSOD-2021-1 to ESK, and the National Institutes of Health under Award Numbers GM118981, GM118982 and GM118983 to HED. Thanks to Dr. Joshua Thomson for many helpful discussions about this work. Contributions: JMF contributed to design, data acquisition and interpretation, and revised the manuscript. HBE contributed to data acquisition and interpretation. CS contributed to data acquisition and interpretation. LC contributed to data acquisition and interpretation. SAA contributed to data acquisition and interpretation. NAP contributed to data acquisition and interpretation. ESK contributed to conception, design, data acquisition and interpretation, drafted and critically revised the manuscript, and performed statistical analyses. All authors give their final approval and agree to be accountable for all aspects of the work. The authors report no conflicts of interest.

## Abbreviations

ETC: electron transport chain

