## Supplemental Figures for "LutABC of *Veillonella parvula* Deacidifies *Streptococcus mutans* Biofilms and Improves Biofilm Health"

Appendix Figure 1

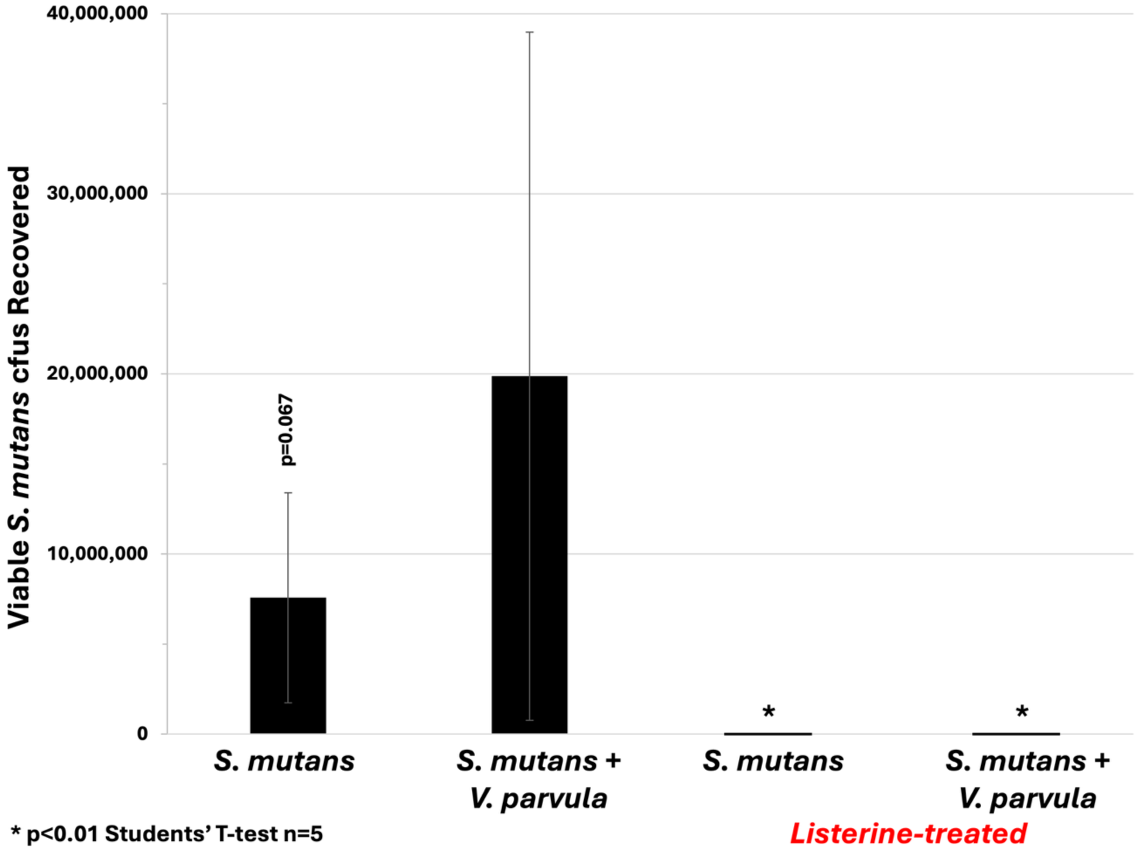


**Appendix Figure 1: *S. mutans* biofilms are viable after 24 hrs in artificial saliva + 0.5% sucrose, even though they stain RED with propidium iodide.** Addition of *V. parvula* leads to a healthier biofilm and shows a trend towards increased *S. mutans* cfu (colony forming units), although the increase does not reach statistical significance ($p=0.067$, $n=5$). Recovered cfu were diluted and plated on BHI agar at 5% $CO_2$ (aerobic) to only permit growth of *S. mutans*. Treatment of either *S. mutans* or *S. mutans* + *V. parvula* for 10 minutes with Listerine naturals prior to plating led to complete loss of viability $>10^6$-fold loss of cfu, 0 cfu recovered.
